# Synaptic correlates of associative fear memory in the lateral amygdala

**DOI:** 10.1101/2020.09.28.317800

**Authors:** Dong Il Choi, Ji-il Kim, Jooyoung Kim, Hoonwon Lee, Ja Eun Choi, Pojeong Park, Hyunsu Jung, Bong-Kiun Kaang

**Author notes:** These authors contributed equally to this work (Dong Il Choi, Ji-il Kim, Jooyoung Kim, Hoonwon Lee).

## Abstract

Successful adaptation to the environment requires accurate responding to external threats by recalling specific memories. However, elucidating underlying neural substrates of associative fear memory was limited due to the difficulties in direct examination of extinction-induced changes of specific synapses that encode an auditory fear memory. Using dual-eGRASP (enhanced Green Fluorescent Protein Reconstitution Across Synaptic Partners), we found that synapses between engram cells or synaptic engram showed a significantly larger spine morphology at auditory cortex (AC) to lateral amygdala (LA) projections after auditory fear conditioning. Fear extinction reversed the enhanced synaptic engram spines while re-conditioning with the same tone and shock restored the size of the synaptic engram. Taken together, we suggest that the synaptic engram may represent a different state of fear memory.

**One Sentence Summary:** Associative fear memory enlarged the spine morphology of synapses between engram neurons in the amygdala, which was diminished by memory extinction and restored by re-conditioning, suggesting that connections between engram cells represent a different state of fear memory.

## Main Text

Successful adaptation to the environment requires making accurate responses to external stimuli, followed by encoding and retrieving these experiences. The behavioral responses should be weakened or enhanced in response to dynamic environments through memory extinction and re-learning processes, and this extinction process facilitates survival by reducing unnecessary energy consumption and refines vital behavioral responses(*1–3*). Recent studies have revealed that engram neurons and synapses are pivotal for memory storage in various brain regions(*4, 5*). In particular, engram tagging and experimental manipulations using promoters of immediate early genes revealed that engram cells are sufficient for retrieval of memory. These innate fear acquisition and extinction processes are well defined in auditory fear conditioning paradigms, which rely on auditory cortex (AC) and auditory thalamus (AT) to lateral amygdala (LA) circuit to establish and recall auditory fear memories(*6, 7*). However, elucidating the underlying synaptic substrates has been constrained without direct examination of specific synapses that encode various states of auditory fear memory. Previous studies primarily utilized electrophysiological approaches that spanned entire inter-regional neuronal connections after fear conditioning and extinction(*8–10*). While the physiological changes induced by fear learning and extinction are well-known, the underlying synaptic correlates that mediate extinction remain unclear. Two mechanisms currently proposed to drive extinction: unlearning(*10–12*) and new learning(*13, 14*) remain elusive in synaptic level as well.

We overcame these challenges by exploiting our recently developed technique, dual-eGRASP, which enables selective labeling of synapses originating from specific neuronal populations, to auditory cortex (AC) to lateral amygdala (LA) circuit (fig. S1A). Dual-eGRASP is an intensified split fluorescent protein, which can only emit fluorescence when pre- and post-synaptic eGRASP are physically attached(*4, 15, 16*) (fig. S1B). By applying dual-eGRASP into AC and AT to LA circuits, which are known to encode auditory fear memory(*6, 7*), we examined whether the synaptic inputs from the two major upstream regions of LA form converged or separated circuits. We found that each LA neuron receives input from both AC and AT (fig. S1C).

We then examined morphological dynamics, such as spine diameter and volume, in two different pathways after fear learning(*17*). To mark the specific synaptic inputs from presynaptic engram neurons, we used a Fos promoter–driven reverse tetracycline–controlled transactivator (rtTA) to express yellow pre-eGRASP in the engram neurons activated during memory acquisition(*18, 19*). We achieved strong expression of cyan pre-eGRASP in a random neuronal population using a high titer of double-floxed inverted open reading frame (DIO) adeno-associated virus (AAV) and a lower titer of Cre recombinase expressing AAV (Fig. 1A)(*4, 20*). After viral infection, we conducted auditory fear conditioning and successfully induced the fear response in both two groups (AC-LA and AT-LA) (Fig. 1, B and C).

**Fig. 1.**
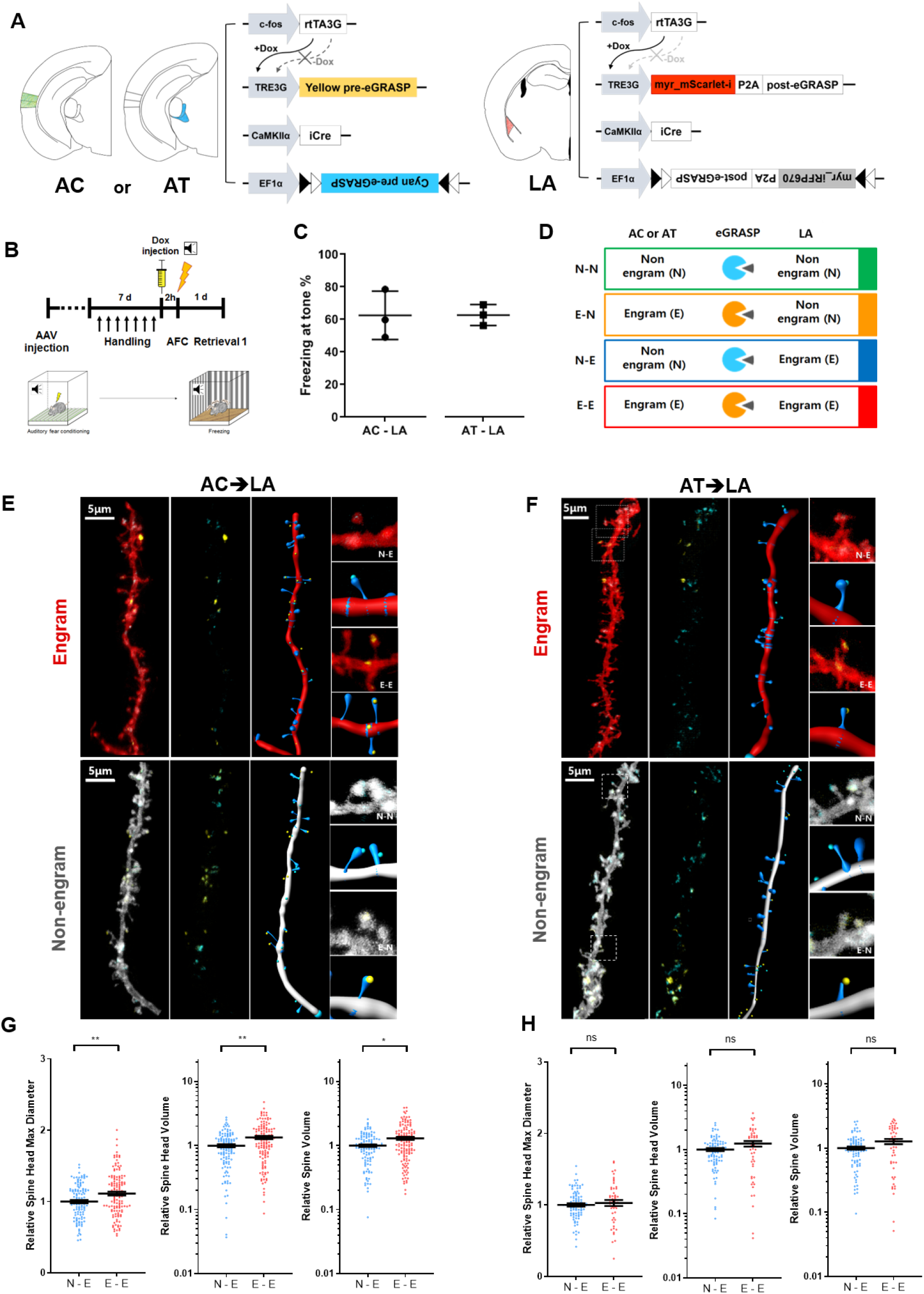
Auditory cortex to lateral amygdala engram synapses exhibited larger spine size after memory formation. **(A)** Schematic illustration of virus injection sites and injected virus combinations of auditory cortex and lateral amygdala. **(B)** Experimental protocol. **(C)** Freezing levels for each group. Each data point represents the freezing levels to tone. AC-LA, n = 3; AT-LA, n = 3. **(D)** Classification of the four synaptic populations indicated by four colors. **(E and F)** Representative images with three-dimensional modeling of lateral amygdala pyramidal neurons for analysis. Red and white filaments indicate engram and non-engram dendrites in lateral amygdala, respectively. Spines from each dendrite are labeled as blue filaments. Yellow and cyan puncta on each spine indicate yellow and cyan GRASP. **(G and H)** Normalized spine head diameter, spine head volume and spine volume on dendrites from neurons. Parameters of spines with yellow puncta were normalized to those of the spines with cyan-only puncta on the same dendrite. n = 116, AC to LA N-E; n = 136, AC to LA E-E; n = 90, AT to LA N-E; n = 50, AT to LA E-E. AC to LA group, n = 3; AT to LA group, n = 3. Mann Whitney two-tailed test, ns = not significant. *p < 0.05, **p < 0.01. Data are represented as mean ± SEM.

All four types of synaptic connections were clearly distinguishable in the same brain slices (Fig. 1D). The spines with only cyan eGRASP signal represent synapses receiving input from non-engram neurons of presynaptic regions whereas the spines with yellow eGRASP signal indicate connections from engram neurons. Engram to engram (E-E) synapses were labeled with yellow eGRASP signals on mScarlet-I-positive dendrites, while non-engram to engram synapses (N-E) were labeled with cyan eGRASP signals on mScarlet-I-positive dendrites. Likewise, engram to non-engram (E-N) and non-engram to non-engram (N-N) synapses were marked by yellow and cyan eGRASP signals on iRFP670-positive dendrites, respectively (Fig. 1, E and F). We measured and analyzed parameters corresponding to spine morphology at each type of synapse in cortico-amygdala (AC-LA) and thalamo-amygdala (AT-LA) circuits. We found a significant increase in spine head diameter and spine volume at E-E synapses in the AC-LA pathway after fear conditioning compared to N-E synapses (Fig. 1G). In contrast, E-N and N-N synapses did not show any significant differences (fig. S2A). These results are consistent with our previous study that showed synapses between engram neurons in CA3 to CA1 circuits are selectively enhanced during fear memory encoding(*4*). However, E-E and E-N synapses in the AT-LA pathway did not show a significant increase of spine size compared to N-E and N-N synapses (Fig. 1H and S2B).

In addition to measuring spine morphology, we investigated the proportion of engram neurons after fear conditioning in AC, AT and LA using Fos-promoter driven nucleus targeting green fluorescent protein (mEmeraldNuc)(*4*). Of note, we found that the engram induction ratio in AT was significantly lower than AC and LA (fig. S3). This result indicates that neurons in AT are less activated by auditory fear conditioning than those in AC and LA.

Next, we investigated whether the enhanced spine morphology could be changed after extinction of auditory fear memory. To distinguish the synaptic combinations in LA, we injected pre- and post-synaptic cocktail virus into AC and LA as same in Figure 1. After viral infection, mice underwent auditory fear conditioning and were divided into extinction and conditioning groups. The conditioning group remained in their homecages, while the extinction group was exposed to the same tone without electric foot shock for three days (Fig. 2A). The series of tone exposures decreased fear response in the extinction group compared to conditioning controls (Fig. 2B). After retrieval 2 for measuring freezing level, we sacrificed the mice and distinguished four possible synapses between AC to LA connections within the same brain slice from both conditioning and extinction groups (Fig. 2C). We measured and analyzed parameters corresponding to spine morphology at each type of synapse (Fig. 2, E to G). Consistent with the results in Figure 1, we found a significant increase in spine head diameter, spine head volume and spine volume at E-E synapses after fear conditioning compared to N-E synapses. In contrast, E-N and N-N synapses did not show any significant differences (fig. S4). Surprisingly, extinction selectively reversed the enhanced spine head size at E-E synapses (Fig. 2, E to G) but did not modify the relative head size of E-N synapses compared to N-N synapses (fig. S4). In addition, as shown in Figure 1, there were no significant changes of spine size between N-E and E-E synapses in the AT-LA circuit in conditioning group (fig. S5). We also did not find any significant differences of spine size between N-E and E-E synapses in the same circuit after extinction (fig. S5). Based on these results, we found that fear conditioning selectively enhanced the size of E-E synapses at least in the AC-LA circuit, while extinction reversed the synaptic enhancement induced by fear conditioning. Therefore, our results suggest that spine head size dynamics form an essential component of memory storage.

**Fig. 2.**
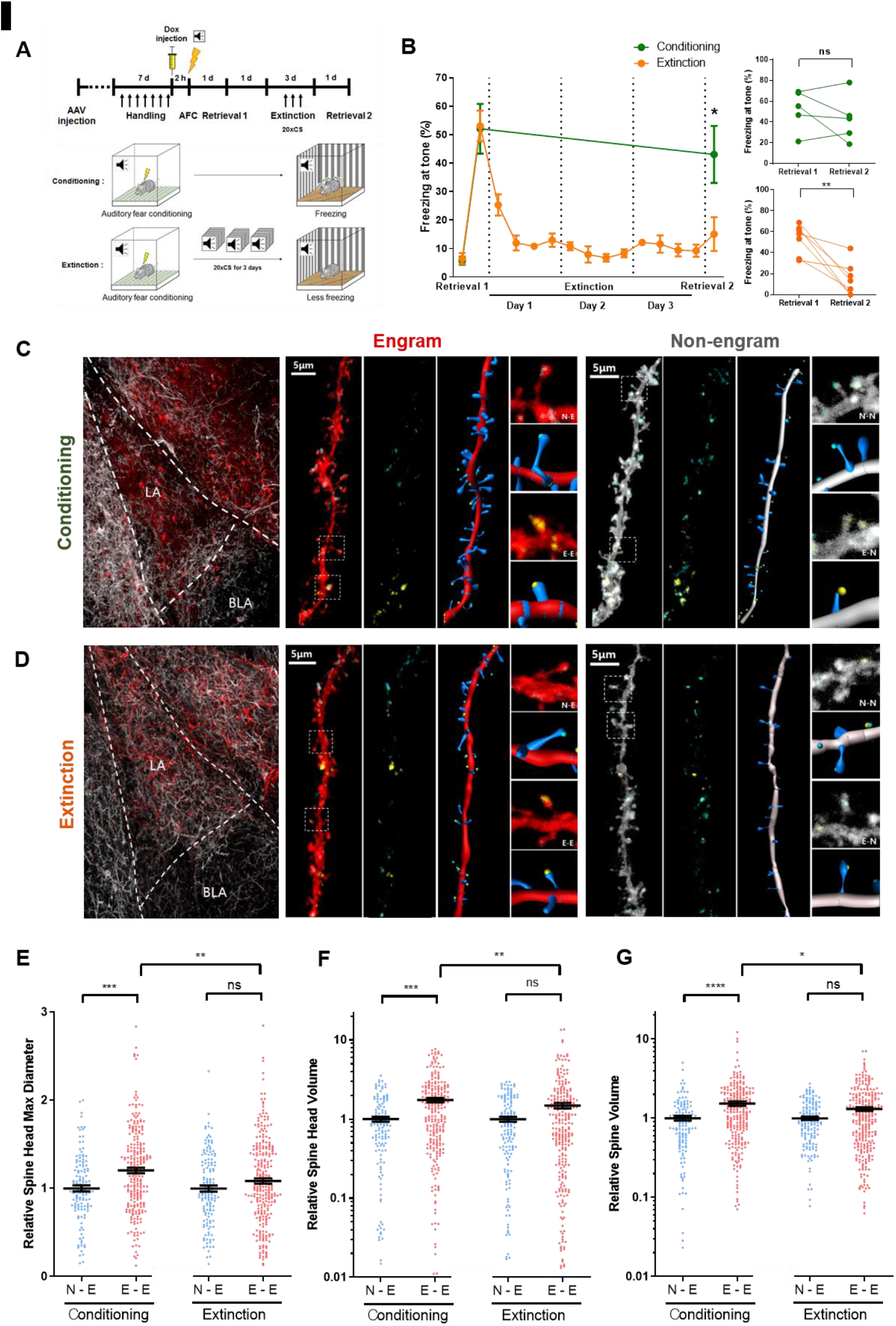
Extinction reversed the fear conditioning-induced enhancement of spine head size at E-E synapses. **(A)** (Upper) Experimental protocol. (Below) Schematic illustrations of the conditioning and extinction processes. Mice were placed into either the conditioning or extinction groups. Both groups were conditioned to an auditory tone. Mice in the extinction group were repeatedly exposed to the tone, while mice in the conditioning group remained in their homecages. **(B)** (Left) Freezing levels for each group. Each data point in extinction session represents the average of freezing levels to tone in 5 minutes. Conditioning, n = 5; Extinction, n = 7; Unpaired t test of freezing levels at retrieval 2; *p < 0.05. (Right, Upper) Freezing levels of mice in the conditioning group were constant across two retrieval episodes at different time points. (Right, Below) Freezing levels of mice in the extinction group decreased after extinction, conditioning, n = 5; extinction, n = 7. Paired t test within each group, ns = not significant, **p < 0.01. **(C and D)**, Representative images of engram and non-engram dendrites with dual-eGRASP labeling in the lateral amygdala of each mouse group. Constitutive cyan dual-eGRASP signals and yellow dual-eGRASP signals expressed by conditioning remained after the extinction process. **(E to G)** Normalized head diameter, head volume, and spine volume on dendrites from neurons. Parameters of spines with yellow puncta were normalized to those of the spines with cyan-only puncta on the same dendrite. n = 126, conditioning N-E; n = 235, conditioning E-E; n = 139, extinction N-E; n = 259, extinction E-E. Conditioning group, mice n = 5; Extinction group, mice n = 7. Mann Whitney two-tailed test, ns = not significant. *p < 0.05, **p < 0.01, ***p < 0.001, ****p < 0.0001. Data are represented as mean ± SEM.

We next speculated that re-conditioning of disappeared fear memory will revive the reduced size of synaptic engram, if the connections between engram cells are an essential component of encoded memory. To investigate this hypothesis, we applied the same viral combination as used in Figure 2. After viral infection, we conducted auditory fear conditioning, following fear extinction behavior. After fear extinction, we divided the mice into extinction and a re-conditioning group. The extinction group remained in their homecages, while the re-conditioning group was exposed to the same tone and electric foot shock for one day as used in the fear conditioning paradigm (Fig. 3A). The freezing response that disappeared by fear extinction was restored by re-conditioning of the fear memory (Fig. 3B). We measured and analyzed parameters corresponding to spine morphology at each type of synapses after retrieval (Fig. 3, C and D). We confirmed no significant difference existed in spine head diameter, spine head volume and spine volume between E-E and N-E synapses in the extinction group, as confirmed in the previous experiment.

**Fig. 3.**
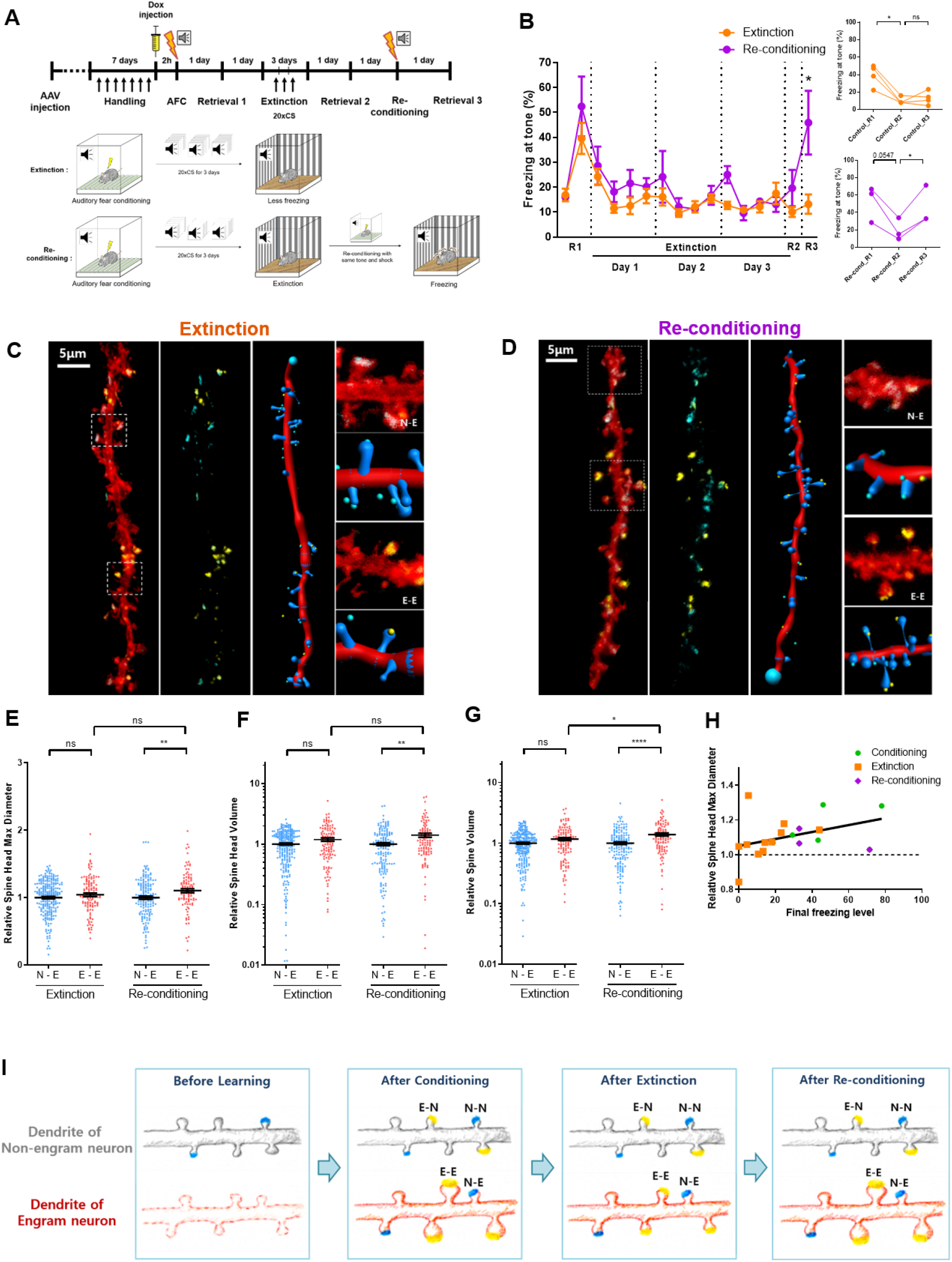
Re-conditioning restored the fear extinction-induced decrease of spine head size at E-E synapses. **(A)** (Upper) Experimental protocol. (Below) Schematic illustrations of the extinction and re-conditioning processes. Mice were placed into either the extinction or re-conditioning process groups. Both groups were conditioned to auditory tone and shock. Mice in the extinction group were repeatedly exposed to the tone, while mice in the re-conditioning group were re-conditioned to same tone and shock after fear extinction. **(B)** (Left) Freezing levels for each group. Each data point in extinction session represents the average of freezing levels to tone in 5 minutes. Extinction, n = 3; Re-conditioning, n = 3; Unpaired t test of freezing levels at retrieval 3; *p < 0.05. (Right, Upper) Freezing levels of mice in the extinction group decreased after extinction at different time points. (Right, Below) Freezing levels of mice in the re-conditioning group were increased after re-conditioning with same tone and shock, extinction, n = 3; re-conditioning, n = 3. Paired t test within each group, ns = not significant, *p < 0.05. **(C and D)**, Representative images of engram and non-engram dendrites with dual-eGRASP labeling in the lateral amygdala of each mouse group. Constitutive cyan dual-eGRASP signals and yellow dual-eGRASP signals expressed by conditioning remained after extinction process. **(E to G)** Normalized spine head diameter, spine head volume and spine volume on dendrites from neurons. Parameters of spines with yellow puncta were normalized to those of the spines with cyan-only puncta on the same dendrite. n = 239, extinction N-E; n = 115, extinction E-E; n = 164, re-conditioning N-E; n = 102, re-conditioning E-E. Extinction group, mice n = 4; re-conditioning group, mice n = 3. Mann Whitney two-tailed test, n.s. = not significant. *p < 0.05, **p < 0.01, ****p < 0.0001. Data are represented as mean ± SEM. **(H)** Comparison between the size of synaptic engram and the freezing level of each group of mice used in Figures 2 and 3. Spine size and freezing level were correlated. Conditioning, n = 4; extinction, n = 11, re-conditioning, n = 3. Spearman correlation test within total mice, *p < 0.05. **(I)** Schematic illustrations representing dynamic changes of each type of synapses among engram and non-engram neurons in the auditory cortex to lateral amygdala circuit by conditioning, extinction and re-conditioning.

Surprisingly, we found that re-conditioning with the same tone and foot shock increased the size of the E-E spine, which was reduced by fear extinction (Fig. 3, E to G). These results indicate that re-conditioning restored the structure of the synaptic engram, which was diminished by fear extinction, rather than creating novel engram synapses. Further, the size of the synaptic engram and magnitude of the freezing level showed positive correlations (Fig. 3H). These results indicate that associative fear memory enlarged the spine morphology of synapses between engram neurons in the AC-LA circuit, which was then diminished by memory extinction and subsequently restored by re-conditioning. Taken together, we suggest that connections between pre- and post-synaptic engram cells or synaptic engrams represent a different state of fear memory (Fig. 3I).

Here, we employed our recently developed synapse labeling technique, dual-eGRASP, to investigate how extinction modified all four kinds of synapses within the AC to LA circuit after inducing an auditory fear memory. Since the enlargement of synapses between engram neurons is the key mechanism underlying memory encoding, we posited these synapses were engram synapses. Previous studies could not identify engram synapses that encode an auditory fear memory in LA given technical limitations. So, structural changes at synaptic engrams after manipulating the fear state, such as extinction and re-conditioning, remained unknown. Based on our results, we conclude that extinction reverses the enhancement at engram synapses and re-conditioning re-increases the diminished engram spine. These data strongly support the unlearning hypothesis of extinction(*10, 12, 21*). However, we cannot exclude that new learning can occur independently through a parallel process. The medial prefrontal cortex may develop an inhibitory circuit with the amygdala during memory extinction(*22, 23*). In addition, previous studies suggested that extinction recruits new engram neurons in the basolateral amygdala after extinction(*23*).

To verify the specificity of the effect, we examined structural changes within the AT-LA circuit. In this circuit, we did not find similar structural changes after conditioning compared to our results in AC-LA circuits, as the population of engram neurons in AT was lower than AC and LA. However, numerous studies provide evidence supporting the idea that both thalamic and cortical inputs into LA are crucial for associative memory formation(*24–27*). Therefore, we cannot exclude other possibilities could underlie our results. For example, synaptic transmission of engram synapses of the AT-LA circuit could be modified after conditioning by other mechanisms, such as changes in the number, type and/or sensitivity of receptors that occur without morphological changes. Further, calretinin-expressing neurons of the lateral thalamus (which includes AT) show plasticity to encode the association of CS and US(*24*). It is possible that the changes of spines between AT and LA may be too subtle to detect, so we cannot conclude that E-E synapses in the AT-LA circuit do not undergo enhancement after conditioning. However, we posit that it is too premature to speculate whether AT-LA synapses utilize different types of synaptic plasticity or whether only specific types of AT cells that may not have been labeled by our dual eGRASP undergo structural synaptic plasticity for unknown reasons.

In this paper, we provided the strong visual evidence of synaptic weakening after fear extinction and its reappearance with re-conditioning. However, demonstrating the causality between E-E spine connectivity and memory such as reduction of engram-specific stimulus-evoked activity is still remained. Overall, our findings demonstrate the physical substrate of memory at the synaptic scale. Our results support the unlearning hypothesis, which postulates that extinction comprises a reversal of conditioning. Future studies will reveal whether our findings are a conserved process for other synapses in the fear learning circuit.

## Acknowledgments

We thank S. Jayakumar and Y. Sung for technical assistance.

## Author contributions

D.I.C, J.-i.K. and B.K.K. designed the experiments. D.I.C., J.-i.K., J.K., H.L., J.E.C., P.P. and H.J. performed the experiments. D.I.C., J.-i.K., J.K. and H.L. analyzed the data. D.I.C., J.-i.K., J.K., H.L. and B.K.K wrote the manuscript. B.K.K. supervised the project.

## Data and materials availability

All data to understand and assess the conclusions of this study are available in the main text or supplementary materials.

## Supplementary Materials

### Materials and Methods

#### Mice

All experiments were performed on 8~10-week-old male C57BL/6N mice purchased from Samtako. Bio. Korea. Mice were raised in 12-hr light/dark cycle in standard laboratory cages and given ad libitum access to food and water. All procedures and animal care were followed the regulation and guidelines of the Institutional Animal Care and Use Committees (IACUC) of Seoul National University.

#### Adeno-Associated virus production

We produced Adeno-Associated Viruses serotype 1/2 (AAV1/2; AAV particle that contains both serotype 1 and 2 capsids) as described in our previous study. Briefly, AAV1/2s were purified from HEK293T cells that were transfected with plasmids containing each expression cassette flaked by AAV2 ITRs, p5E18, p5E18-RXC1 and pAd-ΔF6 and cultured in 18 ml or 8 ml Opti-MEM (Gibco-BRL/Invitrogen, cat# 31985070) in a 150-mm or 100-mm culture dish, respectively. Three to four days after transfection, the medium was collected and centrifuged at 3,000 rpm for 10 min. After 1 ml of heparin-agarose suspension (Sigma, cat# H6508) was loaded onto a poly prep chromatography column (Bio-Rad Laboratories, Inc. cat# 731-1550), the supernatant was loaded onto the column carefully. The column was washed by 4 ml of Buffer 4-150 (150 mM NaCl, pH4 10 mM citrate buffer) and 12 ml of Buffer 4-400 (400 mM NaCl, pH4 10 mM citrate buffer). The virus particles were eluted by 4 ml of Buffer 4-1200 (1.2 M NaCl, pH4 10 mM citrate buffer). The eluted solution was exchanged with PBS and concentrated using an Amicon Ultra-15 centrifugal filter unit (Millipore, cat# UFC910024). The titer was measured using quantitative RT-PCR.

#### Auditory fear conditioning

All mice were fear conditioned 2~4 weeks after the AAV injection. Each mouse was single caged 10 days before conditioning and was habituated to the hands of the investigator and anesthesia chamber without isoflurane for 7 consecutive days. In all experiments, fear conditioning and extinction occurred in two different contexts (context A and context B) to minimize the influence of contextual associations. Context A consist of a square chamber with steel grid floor (Coulbourn instruments; H10-11M-TC), and context B consist of a rectangular plastic box with striped walls and a hardwood laboratory bedding (Beta chip). 2 hours prior to the conditioning, 250 μl of 5 mg/ml Doxycycline solution dissolved in saline was injected intraperitoneally during brief anesthesia by isoflurane. For auditory fear conditioning, mice were placed in the context A and allowed to explore the context for 150 seconds, followed by three exposures to auditory tone CS (30 sec), each of which coterminated with 2 seconds, 0.75mA footshock US, with a 30 sec inter-trial interval^28^. After the conditioning, mice were immediately delivered to their homecages. 1 day after the conditioning, mice were placed into a novel context B and exposed to the auditory tone to measure the freezing behavior. The freezing behavior was recorded and scored using video-based FreezeFrame fear-conditioning system.

#### Fear extinction and re-conditioning

After the auditory fear conditioning, mice were divided into a conditioning group and an extinction group. For three consecutive days, mice in the extinction group were placed into context B. After a 2 minute exploration period, the auditory tone was administered 20 times with a 30 seconds inter-trial interval in the absence of the footshock. Mice in the conditioning group stayed in their homecage during the extinction session. One day after the last extinction session, mice were placed into context B and exposed to the auditory CS to measure the freezing behavior.

For re-conditioning, fear-extinct mice were separated into an extinction group and a re-conditioning group. Mice in the re-conditioning group were re-conditioned under identical conditions as the original auditory fear conditioning procedure. Mice in the extinction group stayed in their homecage during the re-conditioning session. The measurement of freezing behavior was identical to the original procedure.

#### Stereotaxic surgery

Mice (8~10 weeks old) were anaesthetized with a ketamine/xylazine solution and positioned on a stereotaxic apparatus (Stoelting Co.). The virus mixture was injected into target regions using a 32 gauge needle with Hamilton syringe at a rate of 0.125 μl/min. Total injection volume per sites was 0.5 μl, and a tip of the needle was positioned 0.05 mm below the target coordinate right before the injection for 2 minutes. After the injection was completed, the needle stayed in place for extra 7 minutes and was withdrawn slowly. Stereotaxic coordinates for each target sites were: auditory cortex (AP: −2.9/ ML: ±4.5/ DV: −3.2), auditory thalamus (AP: −3.1/ ML: ±1.8/ DV: −3.6), and lateral amygdala (AP: −1.4/ ML: ±3.4/ DV: −4.45).

#### Sample preparation and confocal imaging

Perfused brains were fixed with 4% paraformaldehyde in phosphate buffered saline (PBS) overnight at 4°C, and dehydrated in 30% sucrose in PBS for 2 days at 4°C. Brains were sliced by Cryostat into 50μm section for dual-eGRASP analysis. Sections were mounted in VECTASHIELD mounting medium (Vector Laboratories). For dual-eGRASP analysis, LA dendrites were imaged in Z-stack using a Leica SP8 confocal microscope with 63x objective with distilled water immersion.

#### Image analysis

Processing of confocal image and 3D reconstruction of dendrites were performed using Imaris (Bitplane, Zurich, Switzerland) software. Each mScarlet-I-positive or iRFP670-positive dendrite was manually marked as a filament. Other fluorescent signals were hidden to exclude any bias, and each cyan or yellow eGRASP signal was marked as a cyan or yellow sphere through automatic detection. Overlap of cyan and yellow eGRASP signals were considered as a yellow signal, since the presynaptic neuron of the synapse was c-fos-positive during memory formation. Dendrites without any cyan eGRASP or mScarlet-I, iRFP670-overlapping dendrites were excluded from more precise analysis.

Spines of designated mScarlet-I-positive and iRFP670-positive dendrites were manually reconstructed with automatic detection of diameter and volume. Each spine was defined as an engram or non-engram spine depending on its presence of yellow or cyan eGRASP signal through manual detection. Spine head diameter, spine head volume, and spine length were measured with Imaris FilamentTracer. The examiner was unaware of any eGRASP signals during reconstruction of the spine 3D models.

Normalization of the raw data was done within each dendrite. The raw value of cyan-only spine morphology was averaged, and the raw value of each cyan-only spine and yellow spine was divided by the average. This normalized value was used for further statistical analysis.

#### Statistical analysis

Statistical analyses were performed using Prism 8 (Graphpad). Normality of the data distributions of spine morphology were tested using Shaprio-Wilk test. Comparison of non-normal data sets was tested through Mann-Whitney test.

**Fig. S1.**
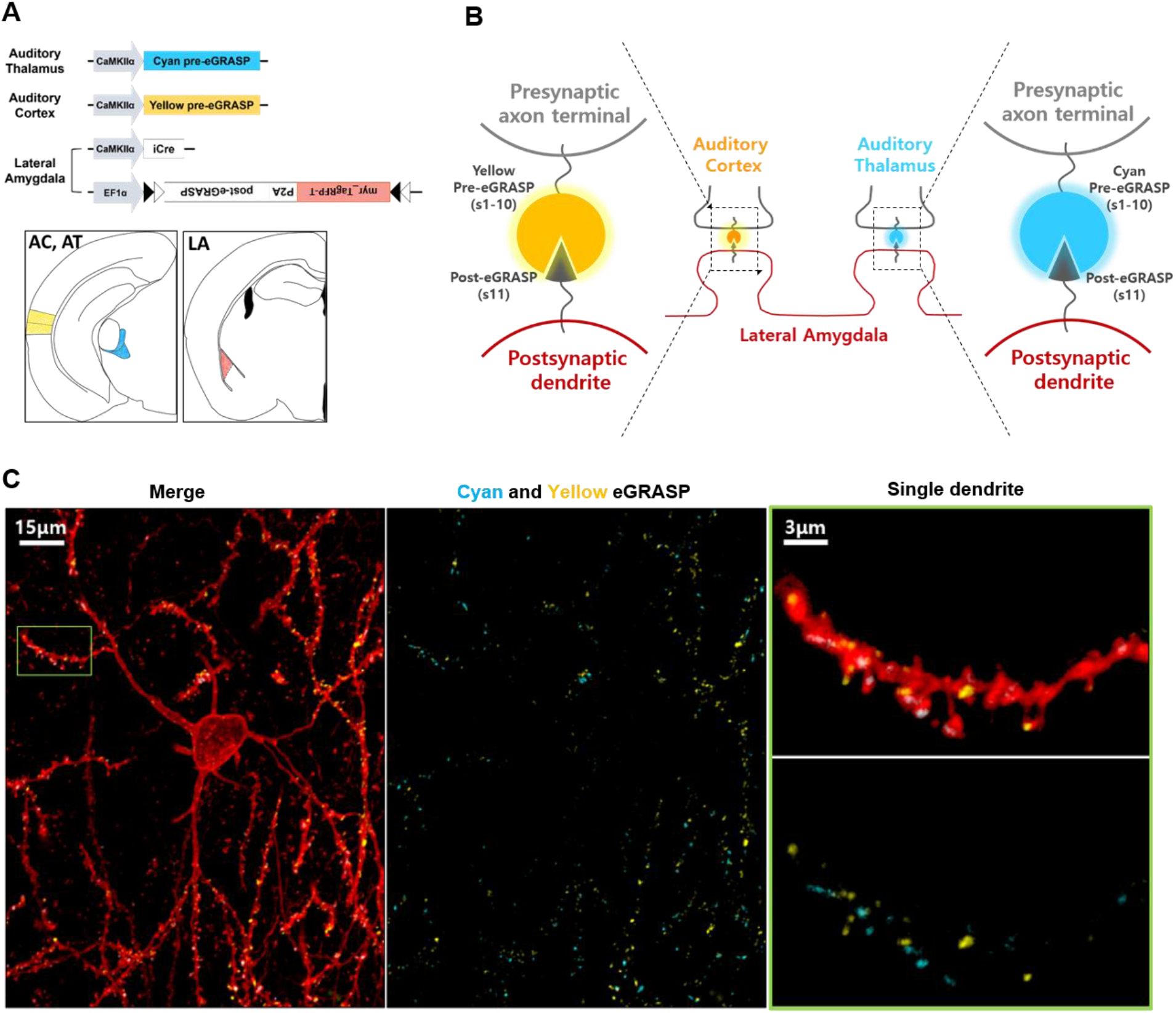
Dual-eGRASP differentially labels synapses on a single lateral amygdala neuron depending on their inputs. **(A)** (Upper) Schematics of injected virus combinations. (Bottom) Illustration of virus injection sites. Each virus containing cyan and yellow pre-eGRASP construct under CaMKII promoter was injected into auditory thalamus and auditory cortex, respectively in one mouse brain. Virus containing myristoylated TagRFP-T (myr_TagRFP-T) and post-eGRASP was injected into lateral amygdala with Cre expressing virus due to sparse expression. **(B)** Illustration of cyan and yellow dual-eGRASP on a single dendrite of a lateral amygdala neuron. Cyan pre-eGRASP and yellow pre-eGRASP were expressed in the auditory thalamus and auditory cortex in the ipsilateral part, respectively. Post-eGRASP was expressed together with myristoylated TagRFP-T (myr_TagRFP-T) in lateral amygdala. **(C)** Representative images of synaptic inputs from auditory thalamus and auditory cortex on a single pyramidal neuron in lateral amygdala. (Left) Post-eGRASP was expressed together with myristoylated TagRFP-T (myr_TagRFP-T). (Middle) Cyan pre-eGRASP and yellow pre-eGRASP were expressed in the auditory thalamus and auditory cortex, respectively. (Right) Enlarged image of single dendrite.

**Fig. S2.**
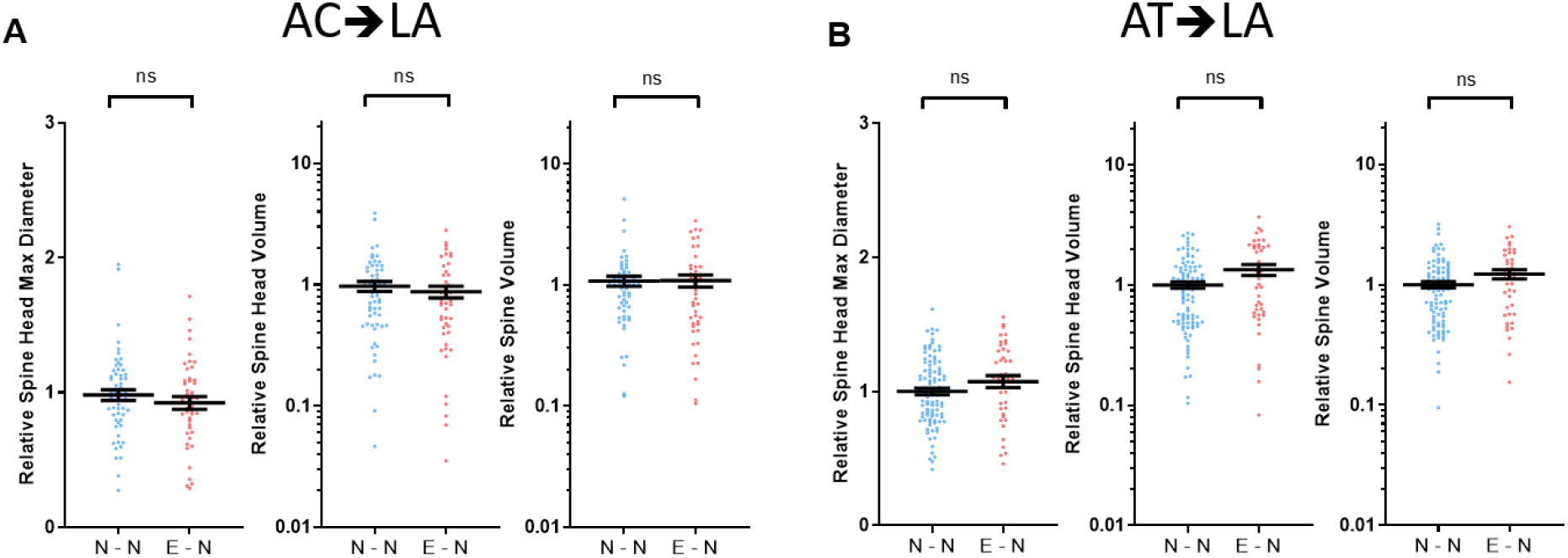
Fear conditioning did not modify the relative head size of non-engram synaptic spine. Normalized spine head diameter and volume on dendrites from neurons. Parameters of spines with yellow puncta were normalized to those of the spines with cyan-only puncta on the same dendrite. n = 59, AC→LA N-N; n = 46, AC→LA E-N; n = 100, AT→LA N-N; n = 42, AT→LA E-N. AC→LA group, n = 3; AT→LA group, n = 3. Mann Whitney two-tailed test, ns = not significant. Data are represented as mean ± SEM.

**Fig. S3.**
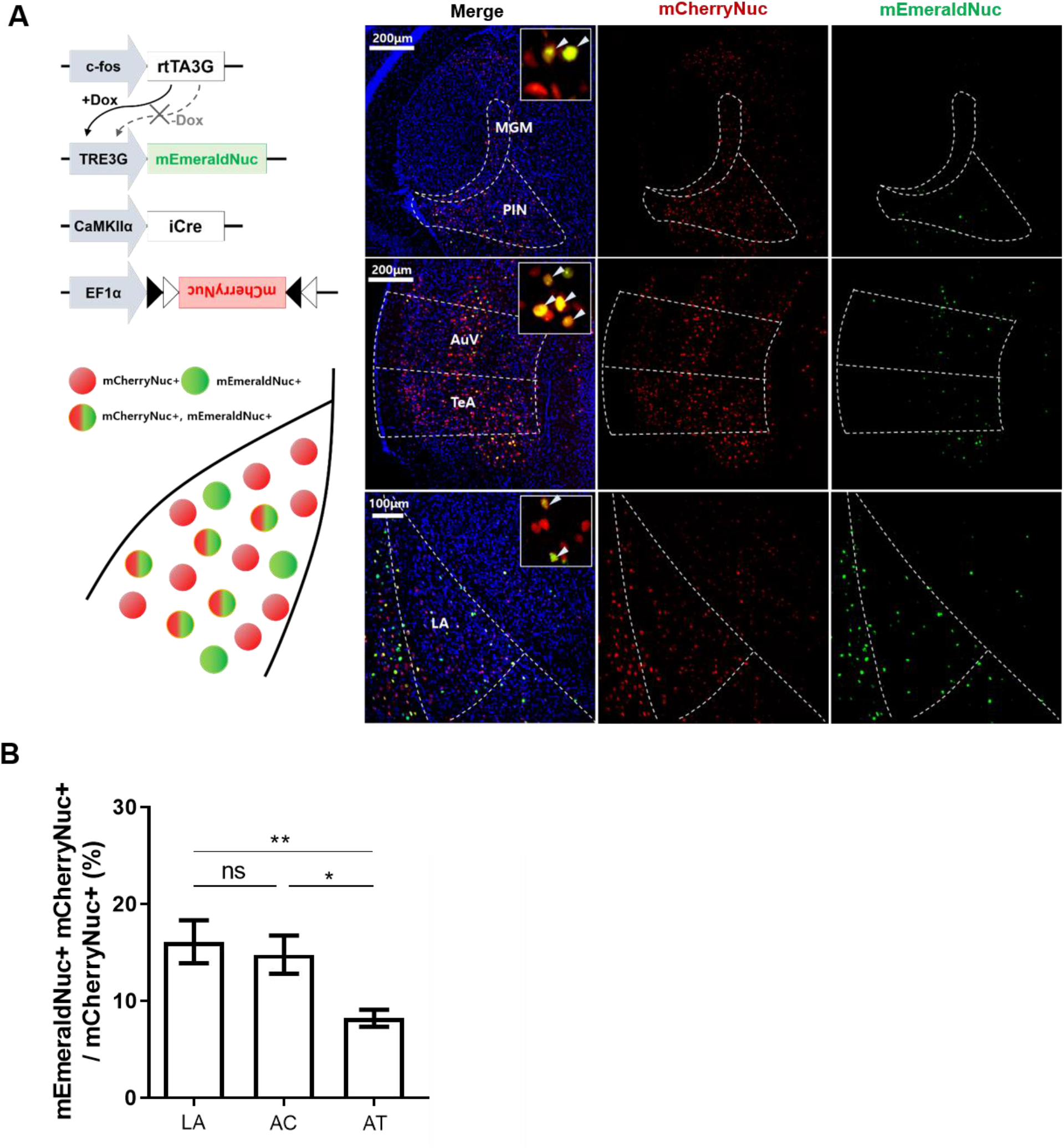
The number of engram labeled neurons was comparable between LA, AC and AT. **(A)** (Left) Schematic illustration of virus combinations and expression pattern of injected site. (Right) Representative confocal images of LA, AC and AT with neuronal labeling by mEmeraldNuc, mCherryNuc. Double-positive neurons were clearly distinguished (arrowhead). AC includes secondary auditory cortex ventral (AuV) and temporal association cortex (TeA), AT includes medial geniculate nucleus (MGM) and posterior intralaminar nucleus (PIN). **(B)** The number of mEmeraldNuc-positive cells in a certain brain region was normalized with a random population of mCherryNuc-positive cells. n = 10, lateral amygdala; n = 10, auditory cortex; n =12, auditory thalamus. Tukey’s multiple comparison test after one-way analysis of variance (ANOVA); *F*(2, 29) = 6.447; n.s., not significant, *P < 0.05, **P < 0.01.

**Fig. S4.**
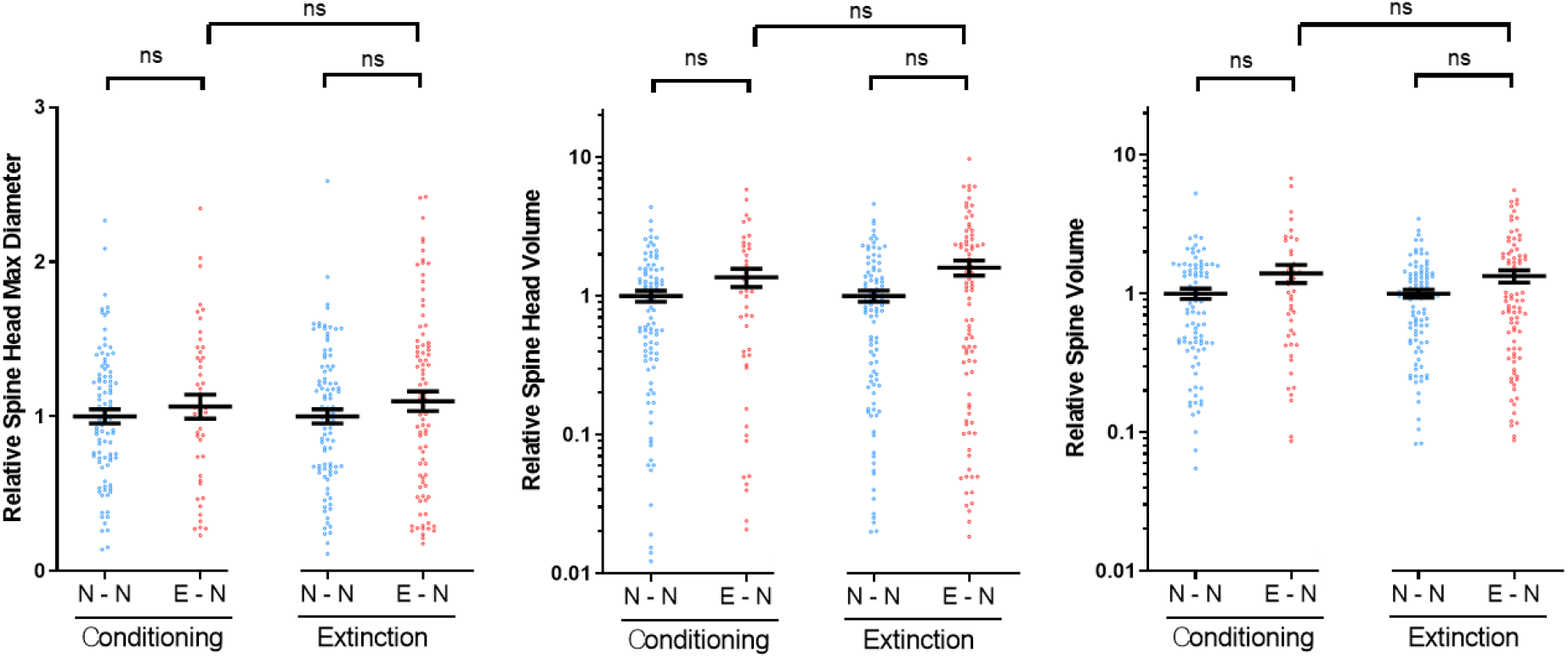
Extinction did not modify the relative head size of non-engram synaptic spine. Normalized spine head diameter, head volume, and spine volume on dendrites from neurons. Parameters of spines with yellow puncta were normalized to those of the spines with cyan-only puncta on the same dendrite. n = 90, conditioning N-N; n = 45, conditioning E-N; n = 97, extinction N-N; n = 87, extinction E-N. Conditioning group, mice n = 3; Extinction group, mice n = 5. Mann Whitney two-tailed test, ns = not significant. Data are represented as mean ± SEM.

**Fig. S5.**
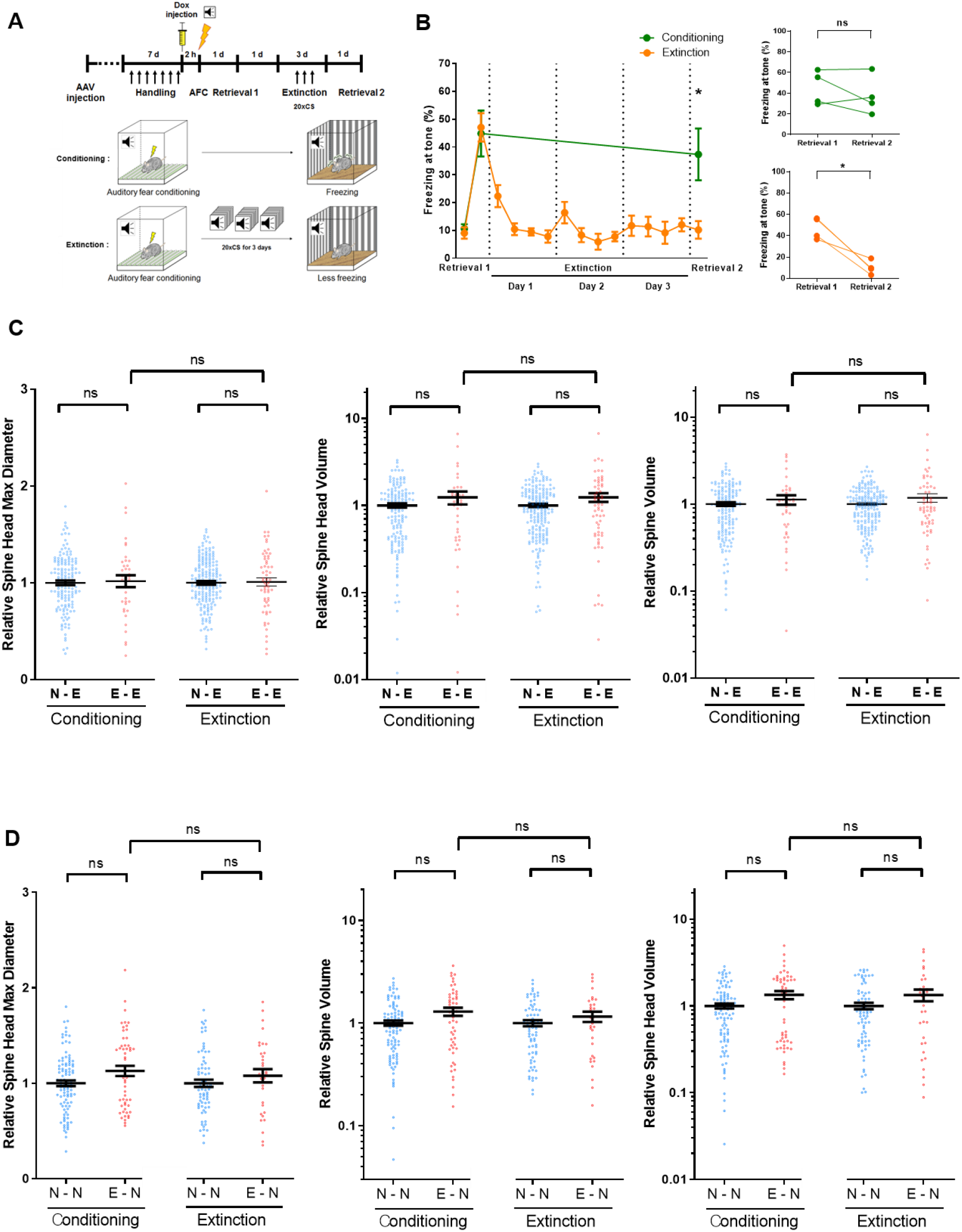
Fear conditioning and extinction did not alter the spine size of thalamo-amygdala synapses. **(A)** (Upper) Experimental protocol. (Below) Schematic illustrations of the conditioning and extinction processes. Mice were placed into either a conditioning or extinction group. Both groups were conditioned to an auditory tone. Mice in the extinction group were repeatedly exposed to the tone, while mice in the conditioning group remained their homecages. **(B)** (Left) Freezing levels for each group. Each data point in extinction session represents the average of freezing levels to tone in a 5 minute period. Conditioning, n = 4; Extinction, n = 3; Unpaired t test of freezing levels at retrieval 2. **(C)** Normalized spine head diameter, head volume, and spine volume on dendrites from engram neurons. Parameters of spines with yellow puncta were normalized to those of the spines with cyan-only puncta on the same dendrite. n = 152, conditioning N-E; n = 38, conditioning E-E; n = 183, extinction N-E; n = 60, extinction E-E. Conditioning group, mice n = 4; Extinction group, mice n = 3. Mann Whitney two-tailed test, ns = not significant. Data are represented as mean ± SEM. **(D)** Normalized spine head diameter, head volume, and spine volume on dendrites from non-engram neurons. Parameters of spines with yellow puncta were normalized to those of the spines with cyan-only puncta on the same dendrite. n = 101, conditioning N-N; n = 54, conditioning E-N; n = 73, extinction N-N; n = 32, extinction E-N. Conditioning group, mice n = 4; Extinction group, mice n = 3. Mann Whitney two-tailed test, ns = not significant. Data are represented as mean ± SEM.

